# Spatial correlations and distribution of competence gene expression in biofilms of *Streptococcus mutans*

**DOI:** 10.1101/2020.11.23.394429

**Authors:** Ivan P. Ishkov, Justin R. Kaspar, Stephen J. Hagen

## Abstract

*Streptococcus mutans* is an important pathogen in the human oral biofilm. It expresses virulent behaviors that are linked to its genetic competence regulon, which is controlled by *comX*. Expression of *comX* is modulated by two diffusible signaling peptides, denoted CSP and XIP, and by other environmental cues such as pH and oxidative stress. The sensitivity of *S. mutans* competence to environmental inputs that may vary on microscopic length scales raises the question of whether the biofilm environment causes spatial clustering of *S. mutans* virulence behaviors, by creating microniches where competence and related phenotypes are concentrated. We have used two-photon microscopy to characterize the spatial distribution of *comX* expression among individual *S. mutans* cells in biofilms. By analyzing correlations in *comX* activity, we test for spatial clustering that may suggest localized, competence microenvironments. Our data indicate that both competence-signaling peptides diffuse efficiently through the biofilm. CSP triggers a Poisson-like, spatially random, *comX* response from a subpopulation of cells that is homogeneously dispersed. XIP elicits a population-wide response. Our data indicate that competence microenvironments if they exist are small enough that the phenotypes of individual cells are not clustered or correlated to any greater extent than occurs in planktonic cultures.

## INTRODUCTION

Oral biofilms are complex microbial communities that may be inhabited by pathogenic as well as commensal species. As a primary etiological agent of dental caries in humans, *Streptococcus mutans* is an important biofilm pathogen (Loesche, 1986; Takahashi and Nyvad, 2011). It forms thick and densely clustered biofilms, especially in the presence of sucrose (Kreth et al., 2004; Leme et al., 2006), consisting of cell clusters embedded in a matrix of insoluble exopolysaccharides, extracellular DNA and other secretions (Watnick and Kolter, 2000; Branda et al., 2005; Xiao and Koo, 2010; Koo et al., 2013). The resulting biofilm is chemically and physically heterogeneous, with parameters such as pH and oxygen concentration varying on microscopic length scales. For example, *S. mutans* fermentation of carbohydrates from the host diet gives rise to localized microenvironments of reduced pH in the oral biofilm (Bowen et al., 2018). At the tooth surface, these acidic niches promote dental caries by demineralizing the tooth enamel (Hunter and Beveridge, 2005; Leme et al., 2006; Takahashi and Nyvad, 2011; Guo et al., 2013; Koo et al., 2013).

In addition to biofilm formation and acid production, the cariogenic behaviors of *S. mutans* include its acid tolerance, carbohydrate utilization, and production of bacteriocins. These behaviors are regulated in connection with the regulation of genetic competence, a transient physiological state in which the microbe can take up and incorporate exogenous DNA (Yoshida and Kuramitsu, 2002; Qi et al., 2004; Ahn et al., 2005, 2006; Senadheera et al., 2005, 2012; van der Ploeg, 2005; Kreth et al., 2006; Welin-Neilands and Svensäter, 2007). The competence pathway in *S. mutans* is sensitive to diffusible signaling peptides as well as to pH, nutrient, and other environmental cues. Because these behaviors are strongly influenced by environmental inputs, one may anticipate that chemical microniches and restricted diffusion within the biofilm shape the spatial distribution of competence gene expression. This raises the question of how virulence behavior is distributed spatially throughout a biofilm, and particularly whether expression is uniformly distributed, or is concentrated or localized into discrete, pathogenic microniches (Stewart, 2003; Parsek and Greenberg, 2005; Hense et al., 2007; Stewart and Franklin, 2008; Bowen et al., 2018).

In streptococci the competent state requires the expression of *comX* (also called *sig*X), which encodes an alternative sigma factor for the late competence genes (Cvitkovitch, 2001; Fontaine et al., 2015; Shanker and Federle, 2017). In *S. mutans,* the activation of *comX* is modulated by inputs such as pH, oxidative stress, and the availability of carbohydrate and peptide nutrients. It is also controlled by two diffusible signaling peptides, denoted XIP (*sigX* inducing peptide) and CSP (competence stimulating peptide). XIP and CSP stimulate the ComRS transcriptional feedback loop through different mechanisms and therefore elicit qualitatively different responses from *comX* in a population of cells (Ahn et al., 2006; Ahn and Burne, 2007; Son et al., 2012, 2015; Guo et al., 2014; Moye et al., 2016; De Furio et al., 2017; Hagen and Son, 2017; Shanker and Federle, 2017).

CSP is produced from its precursor ComC, exported, and then cleaved to length 18 aa. Extracellular CSP interacts with the ComDE two-component system to drive expression of multiple genes associated with bacteriocin production and immunity. Through a mechanism that is not fully known, ComDE stimulates the ComRS system, which is the immediate regulator of *comX*. ComR is an Rgg-like cytosolic regulator that interacts with the 7-residue XIP or its 17-residue precursor ComS to form a transcriptional activator for both *comS* and *comX* (Mashburn-Warren et al., 2010; Shanker and Federle, 2017; Underhill et al., 2019). When CSP is provided to the cell, the ComRS system functions as a noisy, intracellular transcriptional feedback loop. The loop can persist in either the *ON* (high ComS) or *OFF* (low ComS) state but is stimulated by CSP; exogenous CSP therefore triggers a subpopulation of cells (20-40% in planktonic cultures) to switch *ON* and activate *comX* while the rest of the population remains *OFF* (Ferrell, 2002; Son et al., 2012). Bimodal expression of *comX* is a signature property of stimulation by CSP.

By contrast, exogenous XIP can elicit a unimodal response from *comX*: Although *comX* expression in response to XIP varies from cell to cell, it occurs population-wide and is therefore qualitatively unlike the bimodal response to CSP. The unimodal response can be attributed to the import of extracellular XIP (through the Opp permease), which directly interacts with ComR, bypassing the role of endogenous ComS/XIP in triggering the autofeedback loop. Owing to an interaction of growth media with the feedback loop, *comX* responds to XIP only in chemically defined media, whereas it responds to CSP only in complex growth media, rich in small peptides (Hagen and Son, 2017).

The important role of diffusible signals, as well as variables such as pH and oxygen concentration, raises the question of whether *comX* activity in biofilms of *S. mutans* is affected by limited diffusibility within the biofilm matrix, localized chemical gradients, or other physical/chemical factors that are heterogeneous at micron length scales. If activation of *comX* requires exchange of XIP or other signals that diffuse poorly through the matrix or are degraded in transit, then cells activating pathogenic behaviors linked to *comX* could exhibit some tendency to collocate or cluster (Parsek and Greenberg, 2005; Decho et al., 2010). Here we test this hypothesis using two-photon confocal microscopy to probe at single-cell resolution the expression of competence genes throughout the depth of *S. mutans* biofilms. By applying statistical tests derived from spatial ecology we assess the evidence for clustering versus spatial randomness in *comX* active cells. Our data allow us to evaluate the diffusibility of the signaling peptides XIP and CSP and characterize the length scale of any microniches of competence activity.

## METHODS

### Strains

Table 1 shows the UA159-derived strains used in this study. To detect activation of *comX* in *S. mutans* biofilms, and to identify non-expressing cells, we used a dual reporting strain. The *S. mutans* UA159 pGBE-P*comX*::*gfp* / pDL278-P23::DsRed-Express2 strain was constructed using plasmid pDL278-P23::DsRed-Express2, which constitutively drives production of the DsRed-Express2 fluorescent protein encoded on the *E.coli*-streptococcal shuttle vector pDL278 (Shields et al., 2019). The plasmid was transformed into a strain of *S. mutans* UA159 which harbors green fluorescent protein (*gfp*) fused to the promoter region of *comX,* integrated into the chromosome at the *gtfA* site using plasmid pGBE (Son et al., 2018). Plasmid DNA was introduced into *S. mutans* by natural transformation using synthetic CSP peptide (sCSP, see below) synthesized by the Interdisciplinary Center for Biotechnology Research (ICBR) facility at the University of Florida with > 95% purity. The CSP was reconstituted in water to a final concentration of 2 mM and stored in 100 μL aliquots at −20°C.

**Table 1:** Strains and Plasmids used.

| Strain or plasmid | Genotypes and/or descriptions | Source or reference |
| --- | --- | --- |
| ***S. mutans* strains** |  |  |
| UA159 | Wild-type | ATCC 700610 |
| UA159/pDL278 | Wild-type harboring an empty plasmid pDL278, Sp<sup>r</sup> | (Ishkov *et al.*, 2020) |
| UA159/*Pldh-gfp* | UA159 harboring *Pldh-gfp* promoter fusion on pDL278, Sp<sup>r</sup> | (Ishkov *et al.*, 2020) |
| UA159/*PcomX-gfp* (plasmid-based reporter) | UA159 harboring *PcomX-gfp* promoter fusion on pDL278, Sp<sup>r</sup> | (Son, Minjun *et al.*, 2012) |
| UA159/*PcomX-gfp/P23-rfp* (chromosomal *PcomX-gfp* reporter) | Chromosomally integrated *PcomX-gfp* harboring a *P23-DsRed-Express2* red fluorescent protein promoter fusion on pDL278, Sp<sup>r</sup> | This Study |
| UA159/*Pxyl-gfp* | UA159 harboring *Pxyl-gfp* promoter fusion on pZX9, Sp<sup>r</sup> | (Shields *et al.*, 2020) |
| **Plasmid** |  |  |
| pDL278 | *E. coli* – *Streptococcus* shuttle vector, Sp<sup>r</sup> | (LeBlanc *et al.*, 1992) |
| pZX9 | *Streptococcal* replicon with xylose-inducible cassette, Sp<sup>r</sup> | (Xie *et al.*, 2013) |
| *Sp*, spectinomycin |  |  |

For transformations, a final concentration of 2 μM sCSP was added to growing *S. mutans* UA159 pGBE-P*comX::gfp* cultures as they reached OD_600_ nm = 0.2 along with 500 ng of pDL278-P23::DsRed-Express2. After 3 h of additional growth, transformants were plated on BHI agar with the appropriate antibiotics for selection. The strain was then confirmed by PCR using pDL278-specific primers (F - TCA ACT GCC TGG CAC AAT AA; R - TTT GCG CAT TCA CAG TTC TC) and then sequencing of the P23::DsRed-Express2 insert region (Eurofins Genomics). For controls we utilized a constitutively expressing *gfp* reporter under control of the *ldh* promoter as well as UA159 carrying an empty pDL278 plasmid (Ishkov et al., 2020).

The 18-residue form of CSP (SGSLSTFFRLFNRSFTQA) was used throughout. For imaging studies, synthetic CSP, purified to 98%, was provided by NeoBioSci (Cambridge, MA, USA). The rhodamine-B labeled CSP ([Rhod_B] SGSLSTFFRLFNRSFTQA) was synthesized by Pierce Biotechnology (Rockford, IL, USA) and purified to >90%. Synthetic XIP peptide GLDWWSL, corresponding to residues 11–17 of ComS, purified to 96%, was prepared by NeoBioLab (Woburn, MA, USA).

### Growth Conditions

We grew *S. mutans* from glycerol stocks overnight in BBL Brain Heart Infusion (BHI). Strains were grown in an incubator at 37 C and 5% CO_2_ overnight in medium that contained 1 mg ml^−1^ spectinomycin. Cultures were then washed twice by centrifugation and then diluted 25-fold in either complex medium (BHI) or defined medium (FMC) (Terleckyj et al., 1975; De Furio et al., 2017). To facilitate biofilm growth, we supplemented media with either 10 mM sucrose (BHI) or 5 mM sucrose and 15 mM glucose (FMC), both with 1 mg ml^−1^ spectinomycin added. We loaded 300 μl of the prepared culture into a stationary 8 well slide (μ-Slide 8 well glass bottom, Ibidi USA), which was allowed to form a mature biofilm in absence of flow. The Ibidi slide consists of an array of 8 square wells (each 9.4 × 10.7 × 6.8 mm deep) with a glass coverslip bottom that allows the developing biofilm to be observed and imaged with an inverted microscope. Except where stated, the supernatant was exchanged with fresh growth medium after 5 hours of growth. For biofilms that were tested with an inducer peptide, 1 μM CSP or 1 μM (rhodamine B)-CSP (for BHI medium) or 500 nM XIP (for FMC medium) was added. The culture in the 8 well slide was then returned to the incubator for an additional 2 hours before imaging.

### CLSM fluorescence imaging of biofilms

For confocal and multiphoton imaging, biofilms of the *S. mutans* wild-type UA159 and mutant strains were grown in multiwell Ibidi slides. To reduce image noise and enhance green fluorescence, slides were gently washed by removing the growth medium and replacing it with a phosphate buffer solution (Yoshida and Kuramitsu, 2002). Two-photon fluorescence images of the intact biofilms were collected using a Nikon A1RMP Multiphoton Microscope on an Eclipse Ti-E inverted fluorescent microscope frame with a Spectra-Physics Mai-Tai HP DS (Deep See) IR variable (690-1040 nm) pulse laser set to an excitation wavelength of 920 nm. For conventional confocal laser scanning (CLSM) imaging, the excitation source was an LU-N4 Laser Unit set to 488 and 561 nm excitation wavelengths, passed through 525/50 nm and 595/50 nm filter sets respectively. Images were acquired using a 60x water immersion objective (CFI60 Plan-Apo IR, 1.27 NA, WD = 0.16 – 0.18 mm) and each frame was an average of 4 sweeps. Images of the biofilms formed stacks with 1 μm vertical spacing and spanned a total depth of 35 μm. For CLSM imaging of rhodamine-labelled CSP in biofilms of *gfp*-reporting *S. mutans*, a 488 nm excitation laser with a 525/25 green filter was used to collect green fluorescence images and the red fluorescence images were collected using a 561 nm excitation laser and a 595/50 nm filter.

### Image analysis

To probe for structure in the spatial distribution of individual, P*comX*-active cells in the biofilm we evaluated the <u>n</u>earest <u>n</u>eighbor <u>d</u>istance (NND) distribution (Dixon, 2013). The NND distribution *f*(*r*)*dr* is the probability that the distance *r* from a *PcomX-*active cell to its nearest *PcomX-*active neighbor lies in the interval *r* to *r*+*dr*. We found the distribution *f(r)* using a custom Matlab code that evaluates the nearest neighbor distances *r* within one layer of a confocal image stack. We then calculated the <u>c</u>umulative <u>d</u>istribution <u>f</u>unction (CDF) of each NND,

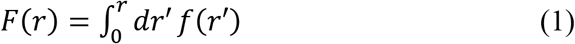

which has the property *F*(*r*) → 1 at large *r*. The experimentally determined CDF can be compared to *F_0_*(*r*), which is the CDF that would occur if an equivalent number of cells were distributed across the image plane by a homogeneous Poisson process, which is often called <u>c</u>omplete <u>s</u>patial <u>r</u>andomness (CSR). If a density *n* (cells/area) of cells is randomly (CSR) distributed over an area, the NND distribution is

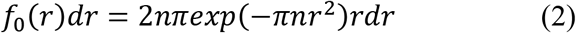

and the CDF is

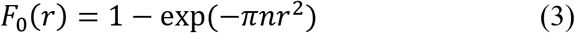

This spatially random CDF must be modified slightly if most cells are randomly distributed, but a fraction *p* are within doublets, or pairs of cells in close proximity (*r* < *r_c_*). For this case we model the expected CDF as

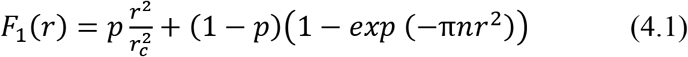

for *r* < *r*_*c*_ and

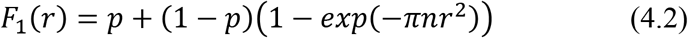

for *r* > *r_c_*. Here *r*_*c*_ is the maximum separation between cells that comprise a doublet.

To assess whether the experimentally observed *F*(*r*) were consistent with the random model *F_1_*(*r*), we (1) fit the CDF obtained from an image showing *N* cells to Equation (4.1) and (4.2) to obtain the parameters (*p, n, r_c_*) for that image, (2) generated a Monte Carlo simulation of *N* cells placed randomly according to those fit parameters, and (3) evaluated the envelope of CDFs from 1000 such simulations. This approach provides a Kolmogorov-Smirnov test of the agreement between the model CDF and the experimental CDF.

In order to assess the randomness of the spatial distribution of P*comX* active cells on distance scales greater than the typical NND, we evaluated the Ripley’s K metric for the distribution of inter-cellular distances (Ripley, 1976; Dixon, 2013). Ripley’s K is widely used in spatial ecology as a test for clustering or dispersion of objects that are distributed in two-dimensions. We used a custom Matlab code to evaluate Ripley’s K for the distances between P*comX*-active cells within an image layer, using the buffer zone method to reduce edge artifacts (Ripley, 1976; Haase, 1995).

## RESULTS

### CSP elicits heterogeneous P*comX* activity in *S. mutans* biofilms in complex medium

CSP elicits a bimodal (subpopulation-only) response from *S. mutans comX* in complex growth media such as BHI, whether cells are grown under uniform, microfluidic flow conditions (Son et al., 2012) or in a biofilm (Aspiras et al., 2004). We first confirmed that our confocal imaging could probe this bimodal response at the individual cell level.

Figure 1 shows two-color CLSM fluorescence images of biofilms of the dual reporting P*comX-gfp*/P*23-rfp* strain growing in BHI medium. As described in *Methods,* biofilms were grown for 5 h in BHI and then incubated for 2 h in fresh medium that contained 1 μM synthetic CSP (Figures 1A, B), or no added CSP (Figures 1C, D), prior to a buffer wash and imaging. The constitutive red fluorescence of individual cells was strong whether CSP was provided or not, revealing the clustered morphology of the biofilms. Green fluorescence indicating *comX* expression was detected only in biofilms that were provided synthetic CSP: Figures 1A, B show P*comX*-active cells as a small, mostly dispersed subpopulation of green cells. This is qualitatively consistent with the bimodal *comX* response to CSP observed in planktonic cultures, although we estimate the *comX-*active subpopulation in the biofilm at only 3-4%, much fewer than the 30-40% of cells that respond to CSP in planktonic conditions (Son et al., 2012; Fontaine et al., 2015).

**Figure 1:**
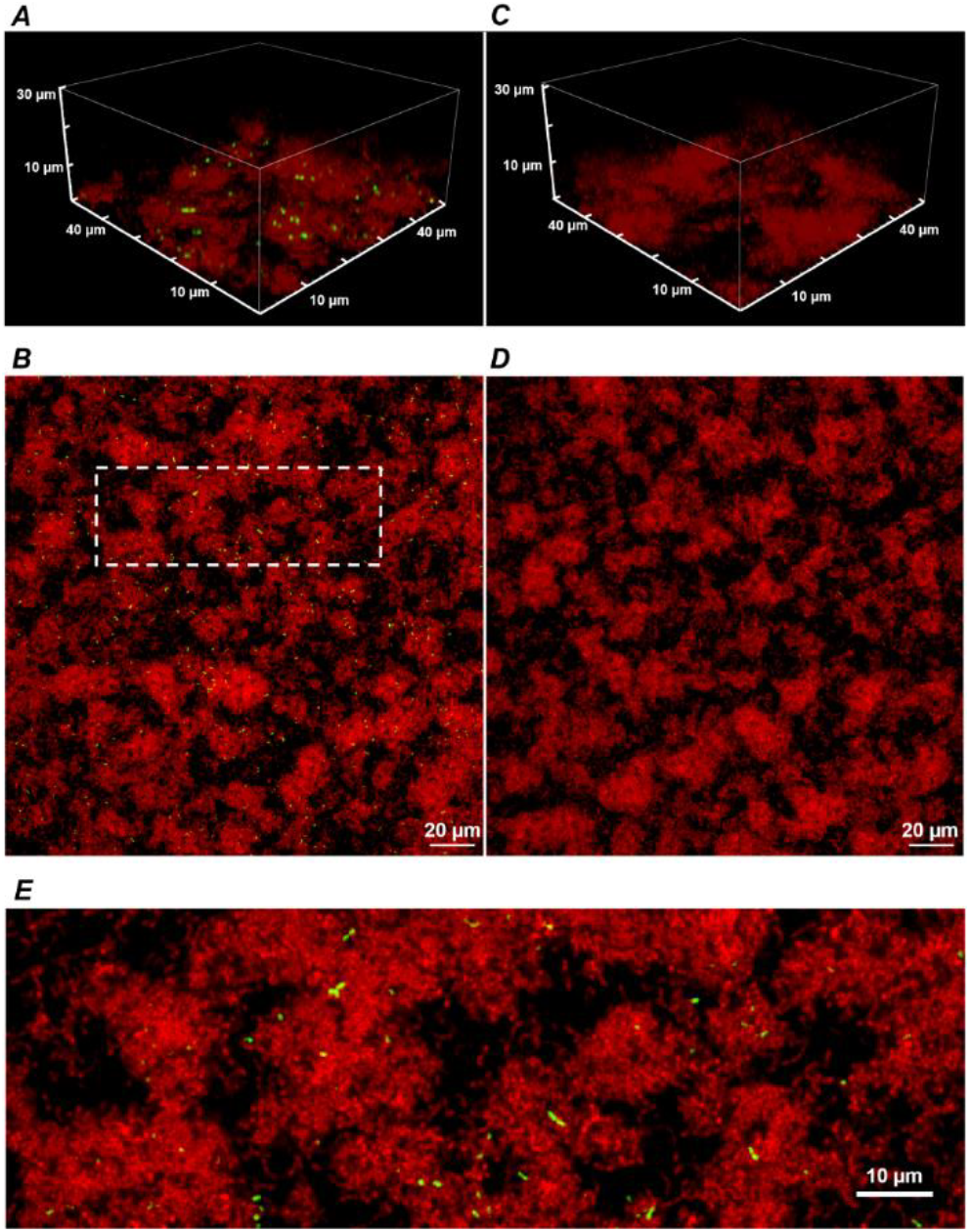
A 3D confocal image stack (**A**) and the substrate layer image (**B**) of a biofilm of the dual reporting (P*comX-gfp*/P*23-rfp*) strain grown for 5 hours before an additional 2-hour incubation in fresh BHI with 1 μM synthetic CSP. Panels (**C**) and (**D**) show the same strain without addition of CSP; (**E**) An expanded view of the dotted region in (**B**). Data shown are representative of 3 separate image stacks collected for each condition.

Supplemental Figure 1 shows the two-color CLSM fluorescence images of biofilms of the non-reporting UA159 strain growing in the same conditions as P*comX-gfp/*P*23-rfp* in Figure 1. As expected, Supplemental Figure S1 shows the absence of any fluorescence on the brightness scale of Figure 1, indicating that the red and green fluorescence in Figure 1 is due to the fluorescent reporters in the P*comX-gfp/*P*23-rfp* strain rather than any background fluorescence.

Although Figures 1B and 1D show that the density of both red fluorescence and P*comX*-active cells was greatest on the bottom (glass contact) layer of the biofilms, individual *comX-* expressing cells are visible throughout the depth of the biofilm and appear mostly well-separated. Figure 1E shows that roughly 10-20% of *comX*-active cells are paired into fluorescent doublets, where each doublet consists of two closely adjacent cells that are axially coaligned to form a bow-tie shape.

### CSP diffuses rapidly through the biofilm

We tested the permeability of an *S. mutans* biofilm to diffusion of CSP. We grew a constitutively reporting P*ldh-gfp* strain in BHI medium with 10 mM sucrose in a multiwell slide. After 5 h, the medium was exchanged for fresh BHI without sucrose and incubated for an additional 4 h. 1 μM synthetic CSP, which was synthesized with an *N*-terminal label of rhodamine B (red) fluorophore, was then added to the different wells at different times. In this way the biofilms were supplied dye-labeled peptide for different time durations, ranging from 7 min to 4 h, before the medium in all wells was exchanged for a phosphate buffer solution and the biofilms were imaged in CLSM.

The red fluorescence of the rhodamine B was seen to quickly concentrate in small pockets of elliptical shape, with length up to 2-3 μm. The pockets are visible throughout the depth of the biofilm, even in the earliest images after 7 min of diffusion. Figures 2A and 2B show these pockets at the bottom (substrate) layer and 5 μm above the bottom, respectively, of each biofilm. This pattern of fluorescence was unchanged after 4 h of diffusion. Interestingly we found no such fluorescent pockets when soluble rhodamine B (no peptide) was added to the biofilm (Supplemental Figure S2), indicating that the pockets of red fluorescence were not due to rhodamine B binding a specific target such as eDNA (Islam et al., 2013). Analysis of the CLSM images (Supplemental Figures S3 and S4) shows that the red fluorescent pockets did not physically overlap with green fluorescence and therefore were not due to staining of live cells. Rather, imaging of planktonic cells incubated with rhodamine-B-CSP (Supplemental Figure S5) indicated that the CSP associates with certain noncellular material. Nevertheless, the rapid permeation of the rhodamine-CSP into the biofilm, with nearly immediate formation of a stable red fluorescence pattern, verified that CSP is mobile through the depth of the BHI-grown biofilm.

**Figure 2:**
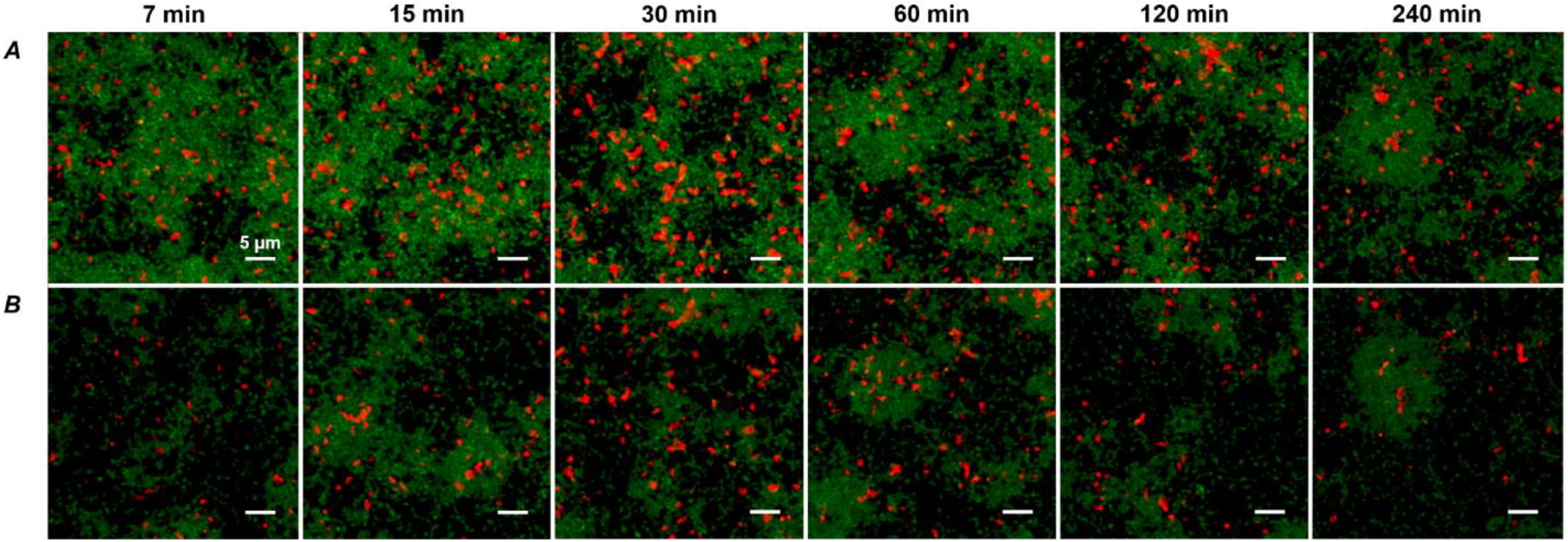
**(A)** CLSM images of the substrate layer of a constitutive *gfp* reporting *S. mutans* strain grown for 5 h in BHI and then incubated (for the time intervals indicated) with 1 μM rhodamine B labeled synthetic CSP. The green channel corresponds to the constitutive *gfp* reporter and the red channel shows rhodamine B fluorescence. The total intensity of the red fluorescence in the image reached its full value within 30 minutes of diffusion. (**B**) CLSM Images at the same *xy* locations as in (**A**) but 5 μm above the substrate. Data shown are representative of 3 separate image stacks collected for each condition.

### *ComX*-expressing cells within a 2D slice of biofilm tend towards a random distribution

We used the distribution of near-neighbor distances (NNDs) as a test of spatial correlations between P*comX*-active cells, which could indicate regions where competence behavior is localized (Dazzo et al., 2013), such as through restricted diffusion of the XIP intercellular signal (Kaspar et al., 2017). Figures 3A, C, E show the distribution *f(r)* of the NND values (distances between nearest neighbors, *Methods*) for P*comX*-active cells in 2D cross-sections such as that of Figure 1B. The image in Figure 1B contains *N =* 492 green cells; the red curve in Figure 3A shows the NND distribution *f*_0_(*r*) that would be expected if 492 cells were distributed across the image plane with complete spatial randomness (CSR, *Methods*). Figure 3B shows *F*(*r)*, the cumulative distribution function (CDF) of the green fluorescent cells, with *F_0_*(*r)* (red curve), which is the NND CDF for a random spatial distribution of 492 cells. Figures 3C-3F show similar analyses at layers further from the substrate. In all cases the true NND distribution and CDF show an excess (relative to the CSR model) at near-neighbor distances near 1-2 μm. This deviation indicates that *comX*-active cells have some tendency to collocate at this separation.

**Figure 3.**
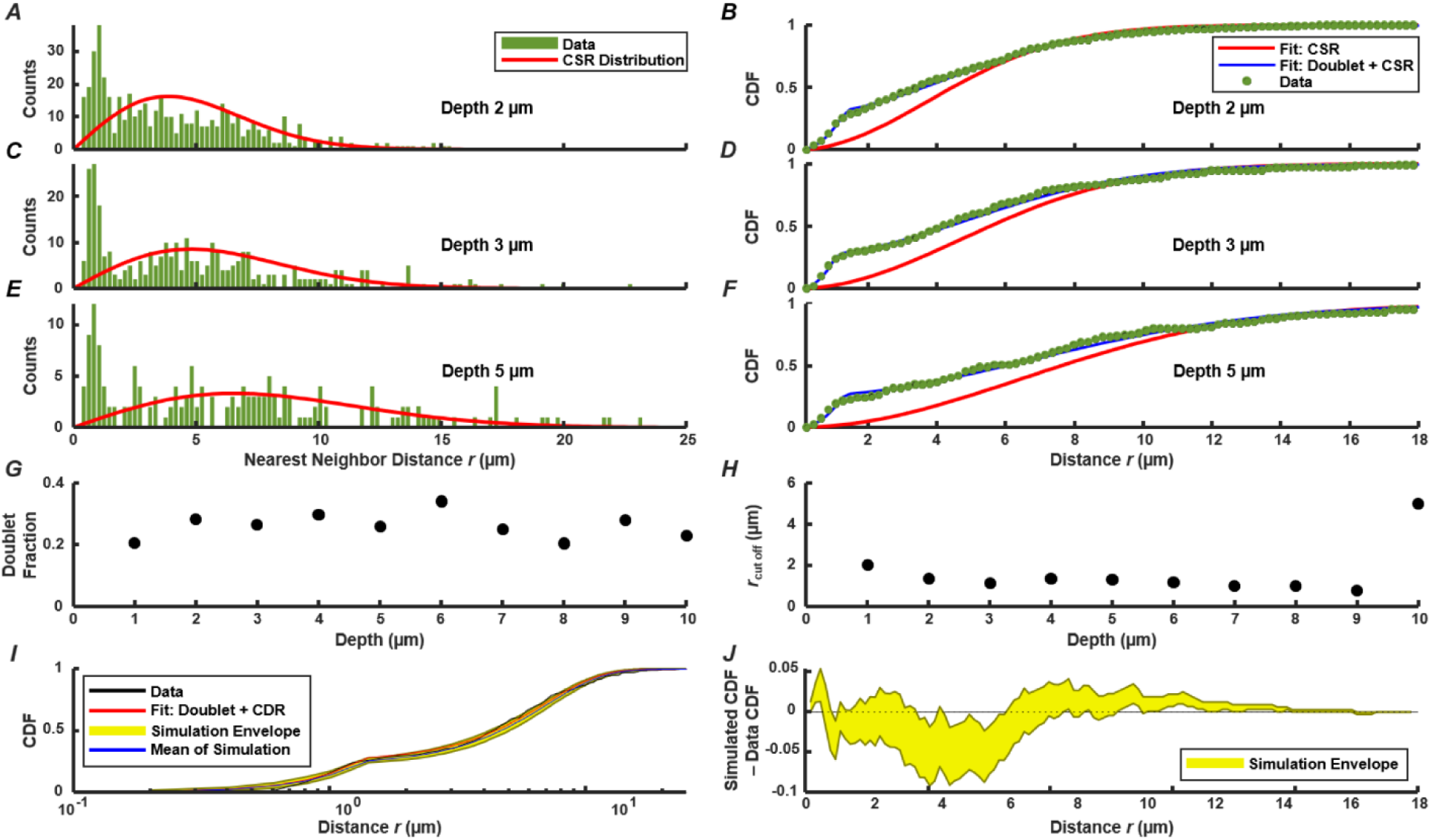
Distribution of nearest neighbor distances between P*comX*-expressing cells in the biofilm of Figure 1A, B following addition of CSP; (**A**) (**C**) (**E**) NND histogram *f*(*r*) (green) of data, and theoretical *f_0_*(*r*) (red) which corresponds to complete spatial randomness (CSR). Panels (**A**), (**C**), (**E**) show *f*(*r*) and *f_0_*(*r*) at depths of 2, 3, and 5 μm respectively within the biofilm; (**B**) (**D**) (**F**) Cumulative distribution function *F*(*r*) of data (green), and theoretical *F_0_*(*r*) (red) which corresponds to CSR. The theoretical *F*_1_(*r*) (blue) for the doublet+random model is also shown in (**B**) (**D**) (**F**) and was obtained from a fit of Equations (4.1) – (4.2) to the experimental *F*(*r*); (**G**) (**H**) doublet fraction *p* and cutoff *r*_*c*_ obtained from fitting experimental CDF to Equations (4.1) – (4.2) at different depths in the biofilm, from 1 to 10 μm above the substrate layer. (**I**) Kolmogorov-Smirnov test by Monte Carlo simulation of the doublet+random model: 1000 simulations were performed of cells randomly distributed in two dimensions with the same total number and density as observed in (**C**), using the fit parameters of *F*_1_(*r*) found in (**D**). (**J**) The difference between the experimental *F*(*r*) and the CDF obtained from the Monte Carlo simulations in (**I**). The yellow envelope in (**I**) (**J**) represents the 90^th^ and 10^th^ percentiles from 1000 simulations.

The confocal images in Figure 1A, B suggest that the anomaly at short distances is a consequence of the cell doublets – pairs of closely adjacent green fluorescent cells. As these doublets have the appearance of sister cells that are not fully separated, we considered that they are likely not due to competence-favoring microenvironments. Rather they correspond to partially attached daughter cells that have inherited the same state of the P*comX* bimodal switch (Hagen and Son, 2017) from their parent cell. Supplementary Figure S6 shows that very similar fluorescent doublets are present after a well-stirred, planktonic culture is incubated with CSP. Therefore we compared our biofilm NND histograms and CDF to a slightly modified spatially random model (*Methods*). The model considers a density *n* of green fluorescent cells that are distributed randomly and homogeneously across the image, where a fraction *p* of these cells have by chance a sister cell within a short distance *r < r_c_*. *F*_1_(*r*) is the CDF for this model. It is given by Equations (4.1) and (4.2), where the cutoff distance *r*_*c*_ defines the maximum separation of cells within a doublet.

Figures 3B, 3D, 3F compare *F*(*r*) (the experimental CDF) with least-squares fits to the doublet model *F*_1_(*r*). The fit parameters *r*_*c*_ ≃ 1.5-2 μm and *p* ≃ 0.2-0.3 are mostly consistent throughout the depth of the biofilm (Figures 3G and 3H). To assess the significance of any disagreement between the experimental CDF and the model we performed a Kolmogorov-Smirnov test, based on a Monte Carlo approach (*Methods*). Briefly, we simulated the random (with doublets) distribution of an equivalent number of points in a two-dimensional plane, using parameters *n*, *r*_*c*_ and *p* obtained from fitting the experimental data. We then evaluated the CDFs for these simulated distributions and compared them to the experimental distribution. As shown by Figure 3I and 3J, the experimental CDF is found to lie within the 10-90% range of the 1000 simulated CDFs. Overall the doublet model provides good agreement with the experimental NND distributions. (Small deviations between the data and Equations (4.1) - (4.2) at distances greater than ~10 μm likely result from finite image size.) The data are therefore consistent with the null hypothesis that, aside from a small number of attached sister cells, the spatial distribution of *comX-* expressing cells in the images lacks structure on length scales less than about 10-20 μm.

To test for spatial correlations on longer length scales, extending beyond nearest P*comX*-active neighbors, we applied a test based on Ripley’s K (Ripley, 1976; Haase, 1995). We calculated, for each P*comX-*active cell in an image, the number of other active cells that lie within a radius *r. K*(*r*) is the average (over all cells) of this number, divided by the mean area density of active cells. By comparing *K*(*r*) to the value expected for CSR (Kiskowski et al., 2009), spatial correlations on long length scales can be detected.

In order to remove the quadratic behavior of *K*(*r*) that is characteristic of CSR, the function *H*(*r*) is often used (Haase, 1995; Hart et al., 2019):

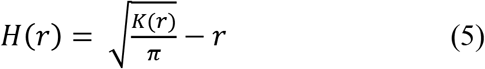

*H*(*r*) = 0 indicates complete spatial randomness, while *H*(*r*)>0 indicates clustering on spatial scales up to *r.*

Figure 4A shows *H*(*r*) calculated for the 492 P*comX*-expressing cells in the 2D layer of Figure 1B. The shaded region is the 90^th^ and 10^th^ percentile envelope from 1000 trials of simulated *H*(*r*), each obtained by evaluating *H(r)* for 492 points distributed randomly (CSR) over equivalent area. The short-range (sister cell) clustering that was detected in the NND analysis causes the experimental *H(r)* to exceed the envelope of the CSR simulations at distances near 2 μm, and *H*(*r*) remains outside the CSR envelope up to distances near 20 μm. Figure 4B shows that including the presence of active sister-cells in the simulation of randomly distributed cells improves the agreement between the experimental and simulated *H*(*r*). However, some possible deviation of *H*(*r*) from the doublet+random model is apparent at *r* = 10-20 μm.

**Figure 4:**
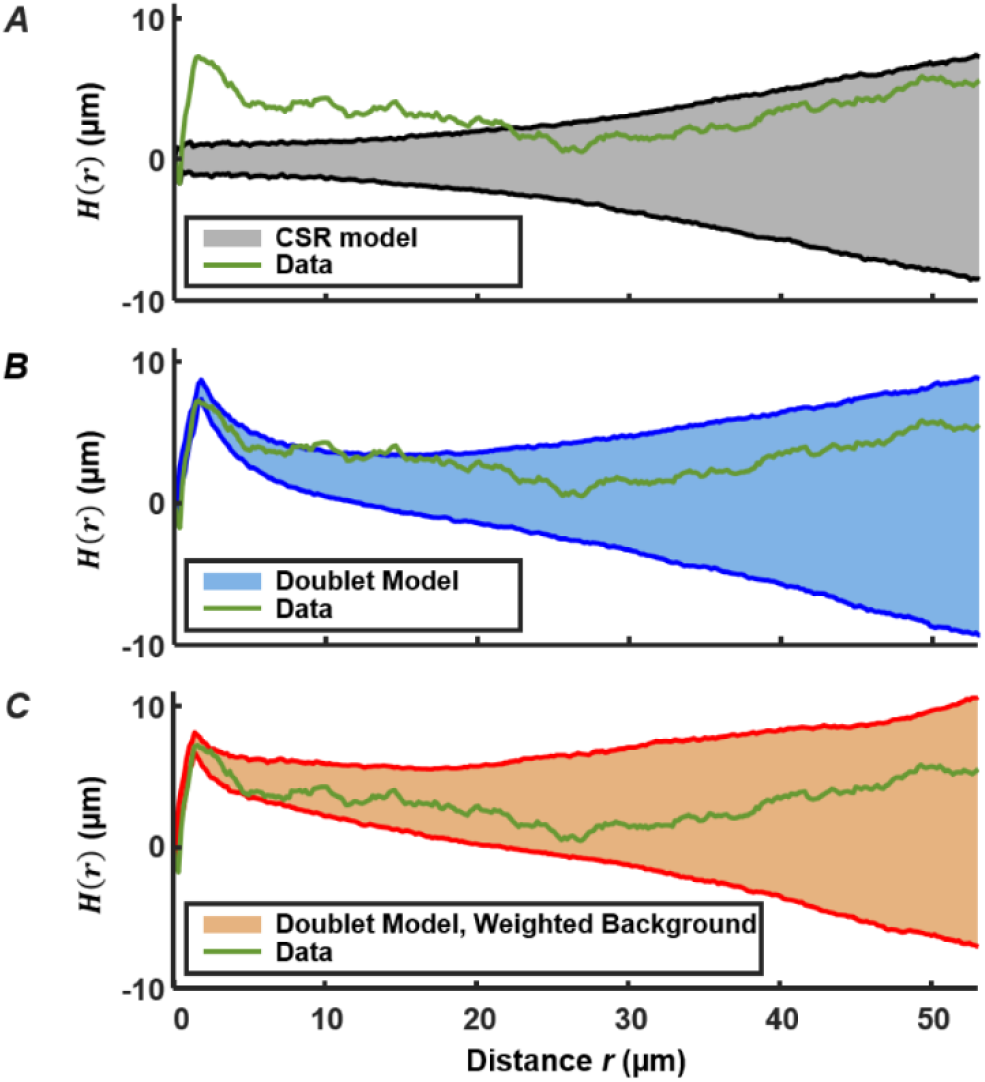
Ripley’s K analysis showing the *H(r)* (green curve) for the *comX*-active cells in the biofilm layer of Figure 1A and 1B. (**A**) Comparison of *H*(*r*) to the envelope of 1000 simulations (shaded region) of *H*(*r*) for 492 cells randomly distributed over the same area. The shading indicates the 10^th^ and 90^th^ percentiles of the simulated *H*(*r*). (**B**) Comparison of *H(r)* to the envelope of simulations of 492 cells distributed by the doublet+random model; (**C**) Comparison of *H(r)* to the envelope of simulations of 492 cells distributed by the doublet+random model, where non-doublet cells are placed at random locations that are weighted according to the red fluorescent (live cell) background of Figure 1B.

We do anticipate some deviation of *H(r)* from CSR behavior on this length scale because the density of the EPS matrix and microcolony biomass itself varies on this scale (Xiao and Koo, 2010). The locations of green fluorescent cells should generally correlate with the distinctly non-uniform pattern of constitutive red fluorescence in Figure 1. Therefore we calculated *H*(*r*) for simulations of a nearly-random spatial model in which (1) a density *n* of green fluorescent cells is distributed across the image, at locations randomly sampled according to a distribution that is weighted by the red fluorescent background in Figure 1B, and (2) a fraction *p* of the green fluorescent cells are part of a sister-cell doublet. This model places green fluorescent cells (including doublets) at random locations, but with a spatial weighting that matches the heterogeneous density of live cells in the biofilm.

Figure 4C compares the experimental *H*(*r*) with that obtained by simulations of this sister-cell + background-weighted model, generated using the *r_c_*, *n* and *p* parameters extracted above. The experimental *H(r*) falls within the 10-90% envelope of 1000 simulations of the model. Consequently the Ripley’s analysis supports the null hypothesis that the distribution of *comX*-expressing cells is generally random and homogeneous, with the exception of sister-cell associations on a very short scale and the variations in the biofilm density on larger length scales. The data give no indication of clustering on scales of roughly 5-20 μm, which might signify microenvironments favorable to *comX* activity.

### Correlation between P*comX* activity and CSP accumulation

Although Figure 2 confirms that CSP permeates the depth of the biofilm, we investigated whether heterogeneity in CSP concentrations on short length scales could affect the spatial distribution of P*comX* activity. We used Figure 2 to test whether *comX* active cells were more likely to lie in proximity to the sites of accumulated CSP revealed by the rhodamine-B label.

Figure 5A shows a confocal image of the substrate layer of a biofilm of the P*comX-gfp* strain, grown in BHI with 10 mM sucrose for 5 h. Spent medium was exchanged for fresh BHI (no added sucrose) containing 1 μM rhodamine-B labeled CSP, and the biofilm was then incubated for 2 h before washing and imaging. Figure 5B shows the histogram of distances from each P*comX*-active cell to the nearest red fluorescent CSP site. Figure 5B also compares that distance histogram to that of a fully randomized distribution, wherein the same number of P*comX-*active cells are placed at completely random locations with respect to the same set of red fluorescence sites. To make that comparison we performed 1000 simulation trials in which the P*comX-*active cells in the image were randomly repositioned within the image field, and the histogram of distances to the nearest red site recalculated; the gray envelope in the figure defines the 90^th^ and 10^th^ percentiles from these simulations. The histogram evaluated from the true locations of the *comX* active cells falls well within this random envelope, indicating the *comX*-expressing cells are not closer, on average, to the CSP-rich sites than would be expected from chance.

**Figure 5:**
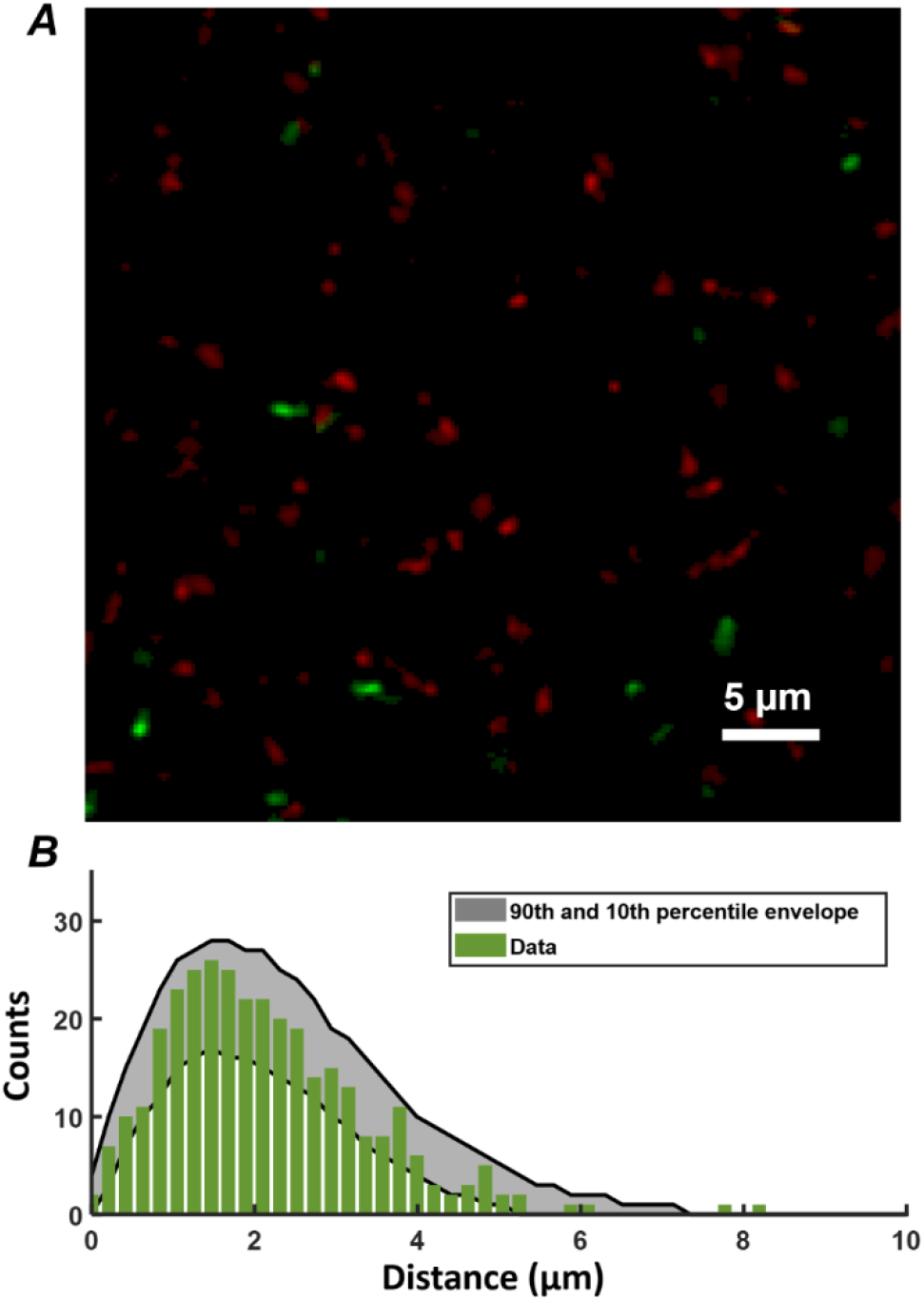
(**A**) Portion of the two-color image of the substrate layer of a P*comX-gfp* (plasmid) biofilm grown for 5 h in BHI and incubated with 1 μM rhodamine-B (red) labeled CSP for 2 h; (**B**) Histogram (green bars) of distances from P*comX*-active (green fluorescent) cells to the nearest rhodamine CSP-rich (red fluorescent) site; The gray envelope represents the 90^th^ and 10^th^ percentiles from 1000 simulations of the green-to-red nearest distance histogram after the *comX-* expressing cells have been randomly relocated throughout the image. Data shown are representative of 3 separate image stacks collected.

### Responsivity to XIP peptide in defined medium declines in aged biofilm

It was previously reported that, when *S. mutans* biofilms were grown in chemically defined medium for periods up to ~20 h, the ability of XIP to induce the (unimodal) *comX* response began to decline (Shields and Burne, 2016). We tested whether this response to XIP remains population-wide (unimodal), as it does for planktonic cells growing in chemically defined medium (Son et al., 2012), or whether the *comX* response occurs within a subpopulation (Shields and Burne, 2016). We grew biofilms of the P*comX-gfp* strain and a xylose-inducible control, P*xyl-gfp*, in the defined medium FMC with 5 mM sucrose and 15 mM glucose for 5, 10 and 20 h. The medium was then exchanged with fresh FMC containing 500 nM synthetic XIP and 15 mM glucose for the P*comX-gfp* biofilms, or 5 mM maltose and 6.66 mM xylose for the P*xyl-gfp* biofilms. Biofilms were then incubated for 3 h and, immediately prior to two-photon imaging, all media were exchanged with phosphate buffer.

Figure 6A-F shows two-photon images of the green fluorescence of biofilms that were grown for 5, 10 or 20 h prior to induction by XIP or xylose. Rather than segmenting individual cells in the confocal image stacks, we characterized the variability in individual cell brightness by coarse-graining the images into cell-sized volumes. We aggregated the data into voxels of 4×4×1 pixels (0.83 × 0.83 × 1 μm^3^= 0.69 μm^3^) and generated histograms of the brightness of these binned voxels.

**Figure 6:**
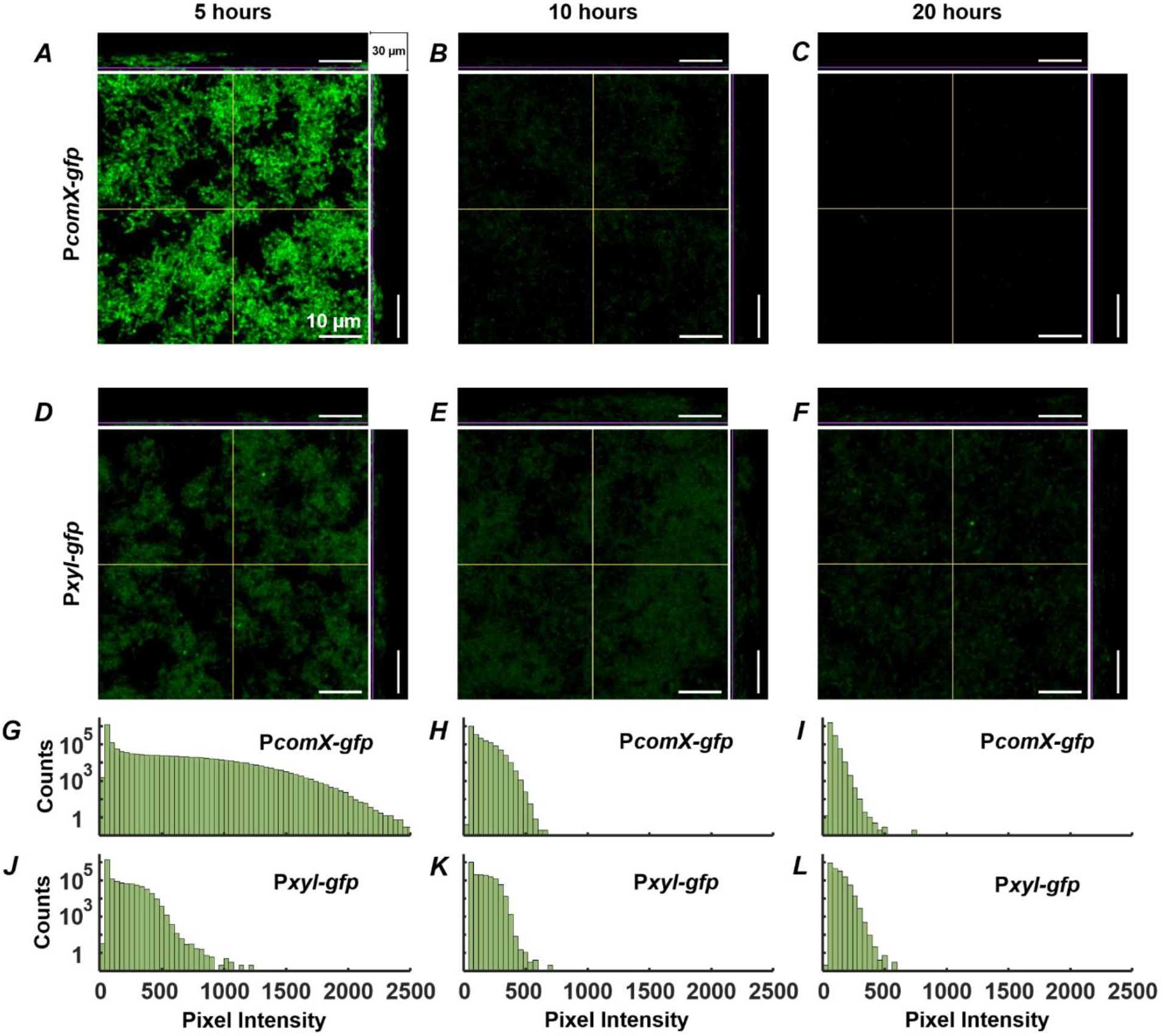
Activation of *PcomX-gfp* (plasmid) grown in FMC media for (**A**) 5, (**B**) 10 and (**C**) 20 h before being incubated with 500 nm XIP for an additional 3 h. A P*xyl-gfp* strain induced with xylose (at the same time as XIP was added for P*comX-gfp*) grown in FMC media for (**D**) 5, (**E**) 10 and (**F**) 20 h under the same conditions as (**A)**-**(C)**; (**G**)–(**L**) Brightness intensity histograms of 16 pixels at a time throughout the depth of the biofilms in (**A**)–(**F**) respectively. All images were obtained by two-photon microscopy with 920 nm excitation wavelength and a 525/25 green filter. The middle panel in each multiphoton image shows the 2D *x-y* plane. The depth (*z*) of this plane within the biofilm is represented by the purple lines on the adjacent top and side panels which show the *x-z* and *y-z* planes respectively. The *x* and *y* regions of the adjacent planes are represented by the yellow lines on the 2D, *x-y* plane. The total stack depth is 30 μm. The brightness scale runs from 0 to 2500 fluorescence units in (**A)-(C)** and 0 to 1500 in (**D)-(F)**. Data shown are representative of 3 separate image stacks collected for each condition.

Both the P*comX-gfp* (Figure 6A) and P*xyl-gfp* (Figure 6D) biofilms show robust fluorescence at 5 h, significantly brighter than the uninduced controls (Supplemental Figure S7). The fluorescence of the P*comX-gfp* is brighter but more heterogeneous than that of the P*xyl-gfp*, although both histograms show unimodal (population wide) fluorescence. The fluorescence of the P*comX-gfp* biofilm declines as the biofilm ages, while the *Pxyl-gfp* biofilm shows a more constant fluorescence histogram over time.

The XIP-induced images are qualitatively unlike the CSP-induced image of Figure 1B, in which isolated, brightly fluorescent cells are separated by substantial regions lacking *comX* activity. The P*comX* response to XIP remains population-wide (unimodal), leading to histograms of GFP fluorescence in Figure 6G-6I that are broad, similar to an exponential distribution, but nevertheless single-peaked. These data indicate that even in the aging biofilm, the mobility of XIP is sufficient, and microenvironments are sufficiently accommodating to competence gene expression, to activate *comX* throughout the biofilm, giving an overall response that is similar to that observed in well-mixed, planktonic cultures.

We note that the P*comX* response to XIP can appear bimodal if the histograms are constructed using a logarithmic fluorescence scale. Supplemental Figure S8 shows that with logarithmic binning, the fluorescence histograms for both the P*comX* and the P*xyl* reporter strains acquire a two-humped character. However, this shape is due to logarithmic binning of a near-exponential distribution; it is not an indicator of bimodal gene expression.

## DISCUSSION

The response of *S. mutans comX* to the exogenous signals XIP and CSP is complex, even when cells are grown in precisely defined, microfluidic conditions (Hagen and Son, 2017). Here we have used confocal microscopy to characterize how heterogeneity in competence gene expression, and particularly the bimodal response of *comX* to the CSP signal, is affected the biofilm environment.

Confocal microscopy has been invaluable for probing the structure of the EPS matrix and the organization of microcolonies in *S. mutans* biofilms (Xiao and Koo, 2010), and for exploring spatial positioning or spatial ecology of *S. mutans* within mixed species biofilms (Nakanishi et al., 2018; Kim et al., 2020). Imaging has shown that cell vitality, respiratory activity and EPS production are distributed nonuniformly in *S. mutans* biofilms (Decker et al., 2014). Several studies have explored competence behavior in biofilms through the imaging of fluorescent gene-reporter strains (Aspiras et al., 2004; Shields and Burne, 2016; Kaspar et al., 2017). Aspiras *et al*. found that green fluorescence in a biofilm of a P*comX-gfp* reporting strain was sparsely expressed following addition of exogenous CSP, with fewer than 1% of cells responding to the stimulus (Aspiras et al., 2004). As both the ComCDE and ComRS elements of the competence pathway have been described as quorum sensing systems, the sparse response of *comX* could imply that restricted diffusion or other local effects in the biofilm modulate expression of the competence genes (Aspiras et al., 2004; Kaspar et al., 2017). The increase of EPS in response to elevated sucrose, for example, could impact the diffusion of XIP within the *S. mutans* biofilm (Klein et al., 2009; Xiao and Koo, 2010; Kaspar et al., 2017). Aspiras *et al.* suggested that *comX* expression may be localized to small clusters of cells, occupying discrete microniches that provide favorable conditions of cell density, nutrient and other factors. Modeling studies have also anticipated spatial correlations or clustering in quorum activity inside bacterial biofilms (Hense et al., 2007; Kindler et al., 2019).

Diffusion in biofilms is affected by molecular charge and mass, interactions with biofilm constituents, and by the porosity of the biofilm (Stewart, 2003; Marcotte et al., 2004; Zhang et al., 2011). High molecular weight solutes such as dextrans and PEG were found to diffuse through *S. mutans* biofilms at half their free (aqueous solution) rate (Zhang et al., 2011), with lower mobility at the base of the biofilm (Marcotte et al., 2004). Uncharged fluorophores like rhodamine B were found to diffuse more slowly than charged fluorophores. However, reductions in diffusion coefficients were generally not large. Although our data suggest that rhodamine B – CSP interacts with a constituent of the matrix, the diffusion coefficient *D* = 1.8×10^−6^ cm^2^/s of rhodamine B in *S. mutans* biofilms was measured at 40% of its value in water, indicating that the biofilm will not dramatically slow the kinetics of equilibration (Zhang et al., 2011). In biofilms of *S. oralis* and *S. gordonii*, even large (40-70 kDa) dextrans penetrated 100 μm cell clusters within roughly 1-2 min (Takenaka et al., 2009).

The above studies predict that the biofilm matrix may have a modest effect on the diffusion of the XIP and CSP peptides. Our data are consistent with that expectation. Although the rhodamine B labeled CSP appeared to concentrate in discrete locations, these locations formed rapidly throughout the depth of the biofilm. In the resulting equilibrium, P*comX*-active cells were not more likely to be near the CSP-rich locations than elsewhere in the biofilm, indicating that their *comX* activity was not triggered by localized high concentrations of CSP. Exogenously-added XIP also appeared to diffuse readily through the biofilms.

Consequently, the spatial distribution of *comX* activity suggests that restricted diffusion does not lead to the formation of clusters or microniches of competence gene expression. The spatial distribution of active cells provides insight into whether other favorable conditions such as pH, nutrient, or other conditions lead to clustering of competence behavior. Dazzo *et al.* demonstrated that statistical methods such as Ripley’s K and the NND distribution (Dazzo, 2012; Dazzo et al., 2013) provide a quantitative and statistically rigorous alternative to methods based on sender/receiver fluorescent reporter constructs (Gantner et al., 2006; Kaspar et al., 2017), and that these statistical tools can be far more sensitive than simple visual inspection of biofilm images. They can identify positive or negative spatial correlations, indicative of cell-to-cell communication. The NND can readily distinguish aggregation from CSR (Dazzo, 2012), while Ripley’s K revealed the presence of two types of local interactions in a biofilm, with length scales of 6-8 μm and 36 μm respectively (Dazzo et al., 2013).

The NND for P*comX*-activity (in response to CSP) was consistent with a simple model in which roughly 80% of *comX*-expressing cells are distributed with spatial randomness – more precisely a homogeneous Poisson process. The remaining 20% of cells are closely proximal to and physically aligned with cells in the first group, indicative of sister cell doublets that have inherited the same state (*ON* or *OFF*) of the *comX* bimodal switch from their parent cell. The excellent agreement between the observed NND distribution and the theoretical distribution for this model supports the null hypothesis that *comX* expression is not significantly clustered into niches or islands. Alternatively, if such niches exist, they are small enough that they comprise at most a solitary cell (or doublet). Although the Ripley’s K analysis indicates some deviations from pure spatial randomness, these effects appear attributable to the correlations between sister-cells and the microcolony morphology of the biofilm itself.

Nevertheless it is interesting that the fraction of cells activating in the biofilm – not more than a few percent - is much smaller than under planktonic conditions. If in the biofilm CSP activated *comX* at the same 30% average rate that is typical of planktonic conditions, the average distance between active cells in the biofilm would have been much smaller in our study. We would then have been able to detect spatial variations in that density, and possibly identify microenvironments in which that activation rate differed from the average. The fact that activation was closer to 1%, which is too sparse to allow any such analysis, suggests that conditions throughout the biofilm are generally unfavorable. Local conditions such as pH could bias the *comX* bimodal switch to the extent that very few cells are able to accumulate enough intracellular ComS to activate the transition from *comX OFF* to *ON.* In this case the competence microniches consist of rare locations where one cell (or a doublet) is able to activate. To test this possibility it may be useful in future studies to apply spatially resolved probes of key environmental inputs such as pH and nutrient condition and correlate these with *comX* activity.

## ACKNOWLEDGMENTS

The authors acknowledge microscopy assistance provided by Doug Smith of the UF Cell and Tissue Analysis Core, as well as discussions with Prof. Robert A. Burne and Dr. Robert C. Shields.

## AUTHOR CONTRIBUTIONS

II designed and performed experiments, analyzed and interpreted data and wrote the manuscript, SH designed experiments, analyzed and interpreted data, and wrote the manuscript. JK interpreted data, constructed bacterial strains and edited the manuscript. All authors gave final approval to the manuscript and agree to be accountable for all aspects of the work.

## CONFILICT OF INTEREST STATEMENT

The authors declare no conflicts of interest.

