## Supplemental Figures for "Spatial correlations and distribution of competence gene expression in biofilms of *Streptococcus mutans*"

### Supplemental Information

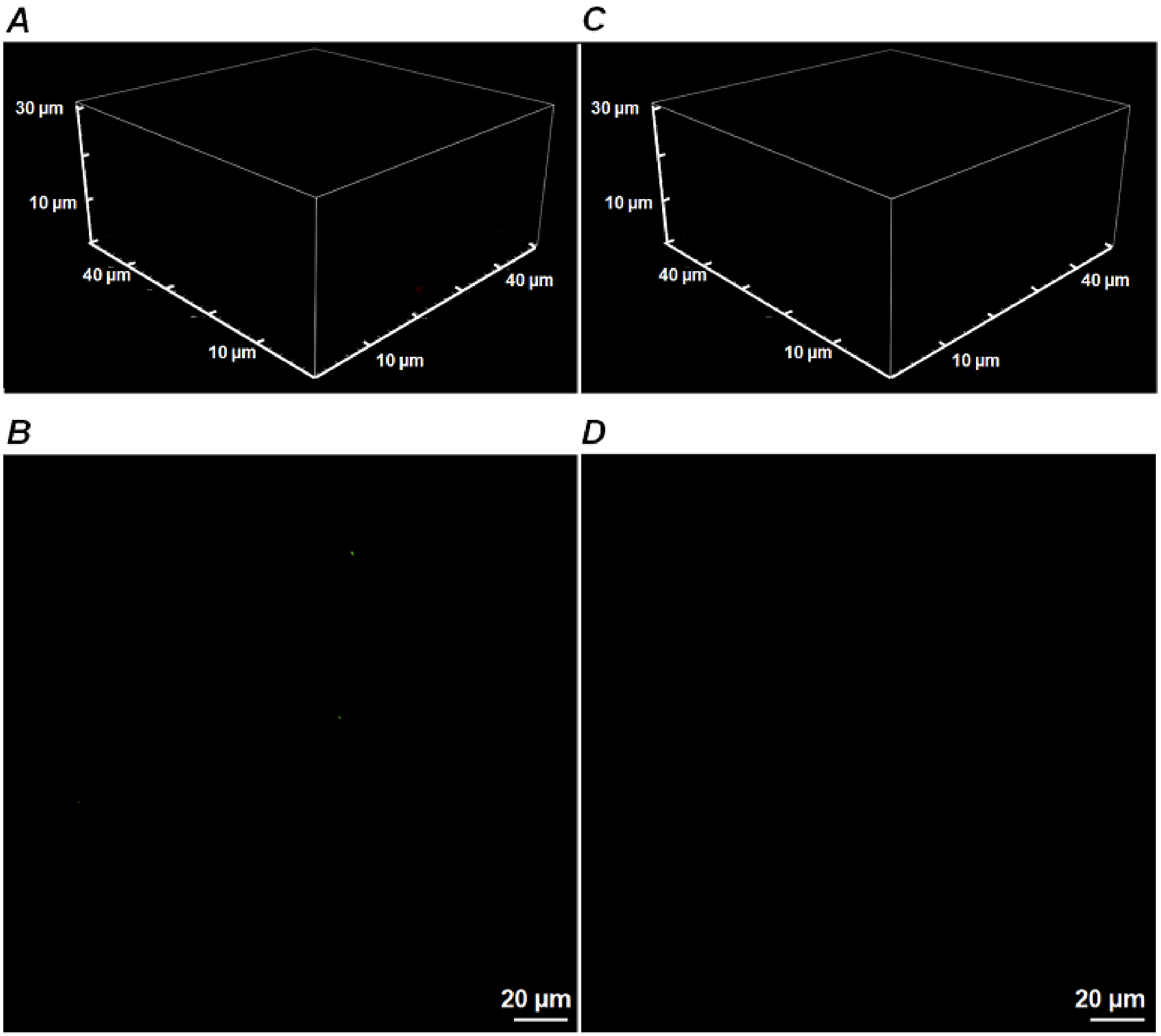

**Supplementary Figure S1**: UA159 fluorescence controls show no green or red fluorescence for growth in BHI +/- CSP.

CLSM images of green and red fluorescence of biofilms showing absence of any fluorescence in the background strain UA159 (no *gfp* or *rfp* reporter): (A) 3D image and (B) bottom (substrate) layer green and red fluorescence image of UA159 grown for 5 hours, followed by an additional 2-hour incubation in fresh BHI with 1 µM added synthetic CSP; (C) 3D and (D) bottom layer green and red fluorescence image of UA159 prepared under identical conditions without addition of CSP. Image brightness scale is the same as in Figure 1. Data shown are representative of 3 separate image stacks collected for each condition.

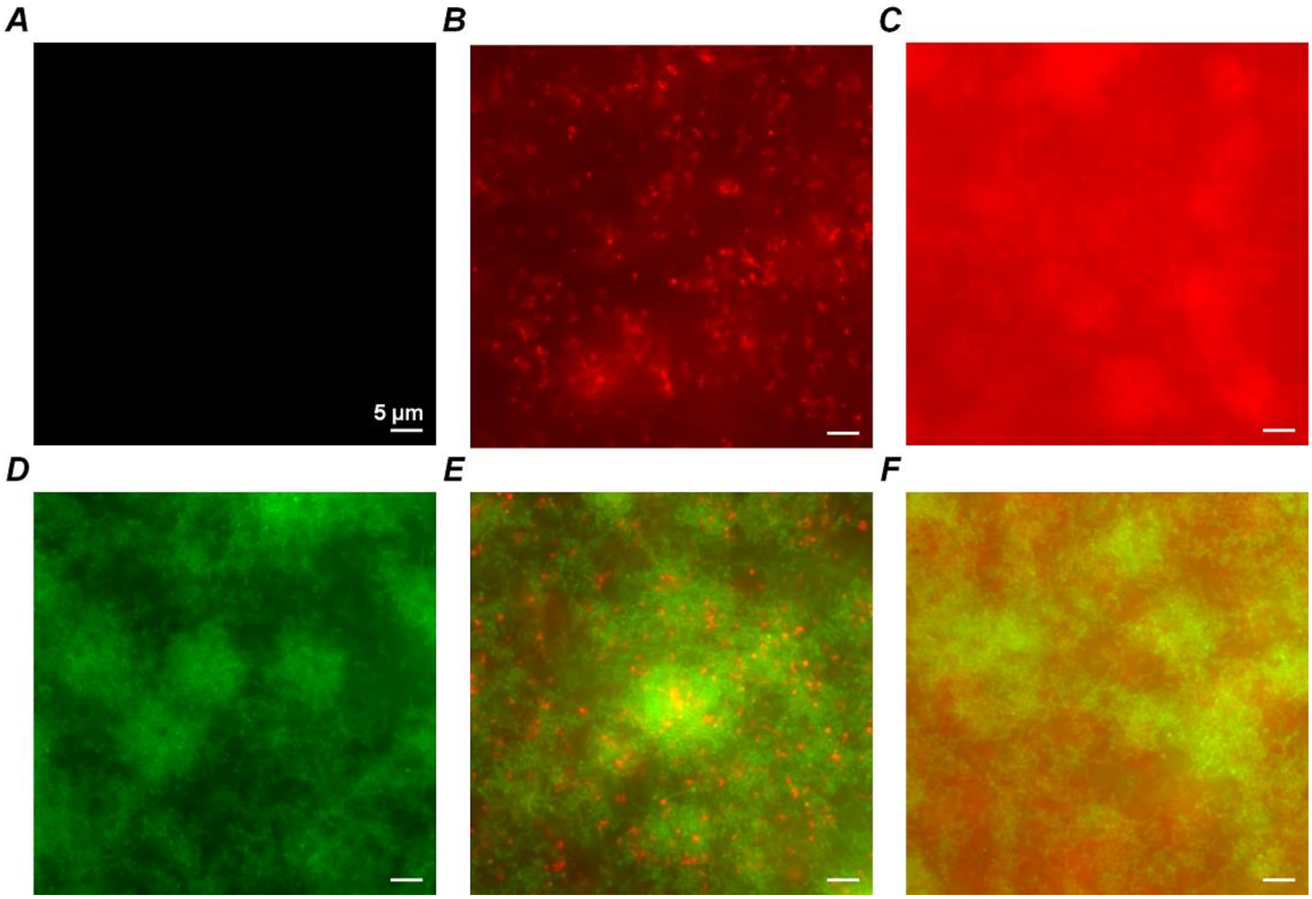

**Supplementary Figure S2**: Comparing diffusion rhodamine B and rhodamine-B labeled CSP within the biofilm of *S. mutans*.

Fluorescence microscopy images of *S. mutans* biofilms of (A)-(C) UA159 background and (D)-(F) P*ldh-gfp* reporting strain. Green (GFP) and red (rhodamine-B) fluorescence images are overlaid. Biofilms were grown for 5 h in complex (BHI) medium before being incubated for an additional 2 h in fresh BHI containing either (A),(D) no rhodamine, or (B),(E) 1 µM rhodamine B labeled CSP, or (C),(F) 1 µM rhodamine B (no CSP). The rhodamine fluorophore by itself permeates the biofilm without showing any visible aggregates; however, when the fluorophore is attached to the CSP the biofilm shows points of concentrated red fluorescence similar to those observed in confocal microscopy, Figure 2. Images were obtained using a Nikon TE2000U phase contrast / fluorescence inverted microscope. Data shown are representative of 3 separate images collected for each condition.

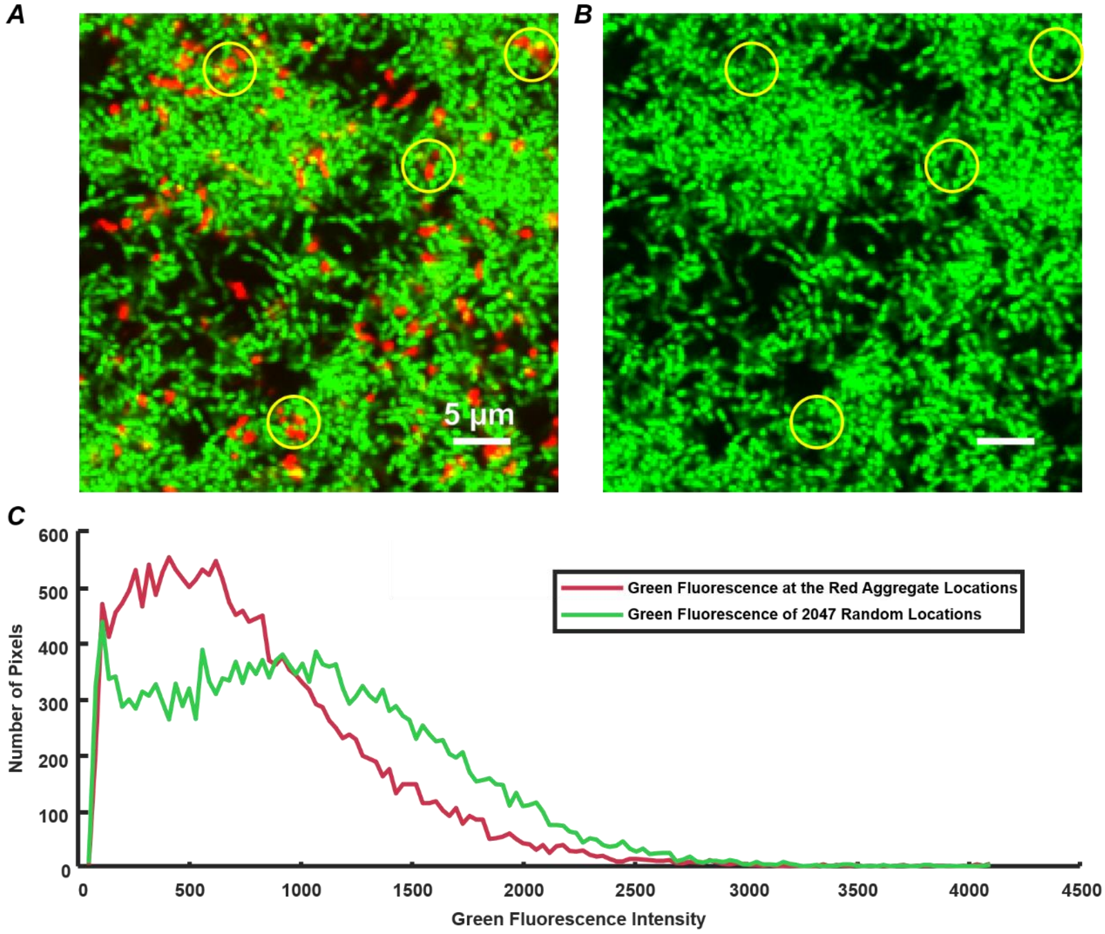

**Supplementary Figure S3:** Comparing location of rhodamine B labeled CSP accumulation and *gfp*-expressing *S. mutans* cells in biofilm

Analysis of red and green fluorescence locations in CLSM images of Figure 2. (A) CLSM images of the substrate layer of a constitutive *gfp* reporting *S. mutans* biofilm grown for 5 h in BHI and then incubated for 30 min with 1 µM rhodamine B labeled CSP. The green channel corresponds to the constitutive *gfp* reporter and the red channel corresponds to rhodamine B red fluorescence; (B) Green channel of (A) without the red channel overlay, with yellow circles indicating locations of selected rhodamine B labeled CSP accumulations in (A); (C) histograms of green fluorescence intensity for two different samplings of locations in (A): histograms of green fluorescence at 2047 locations where red fluorescence is observed (red curve), and histogram of green fluorescence at 2047 randomly selected locations that lack red fluorescence (green curve). To reduce the effect of noise in the comparison, the resolution of the original image (A) was reduced by 3x3 pixel binning.

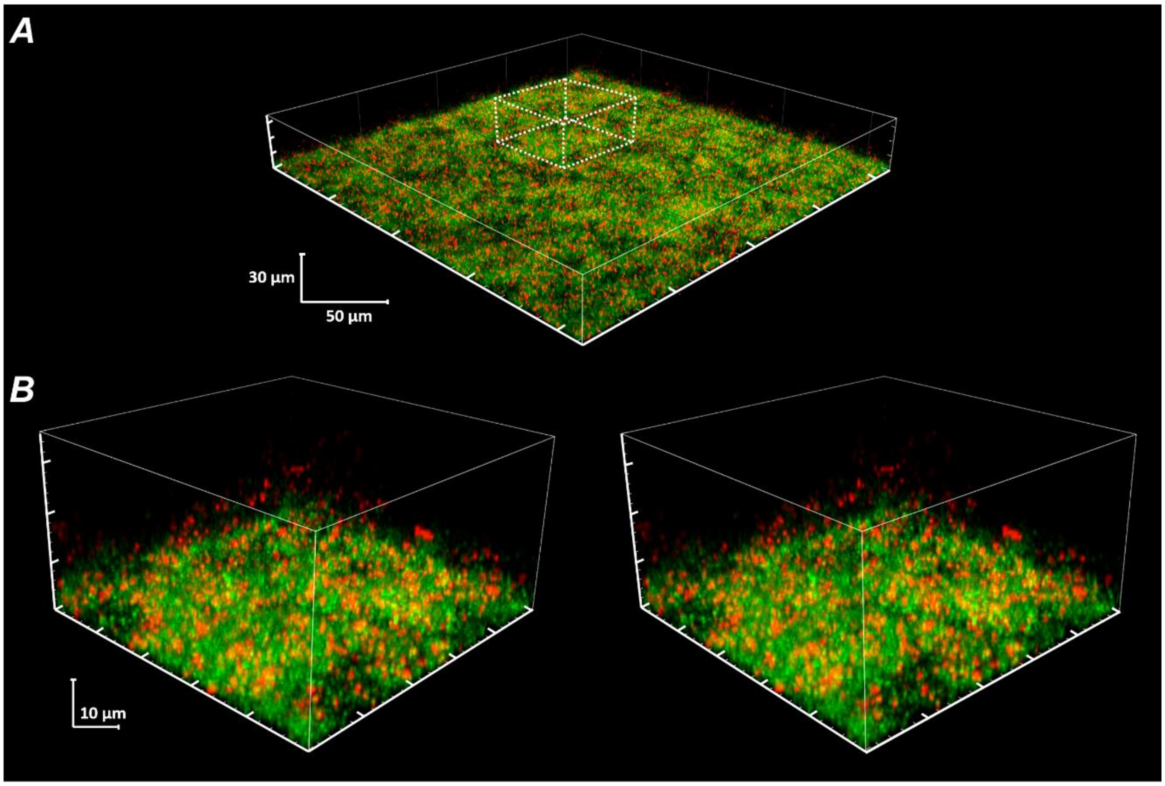

**Supplementary Figure S4:** Three-dimensional spatial distribution of rhodamine B labeled CSP within a *gfp*-expressing *S. mutans* biofilm.

(A) CLSM image of a constitutive *gfp* reporting *S. mutans* strain grown for 5 hours in BHI and then incubated with 1 µM rhodamine B labeled CSP for 30 mins in fresh BHI. The red channel (561 nm excitation with a 595/50 filter) shows the distribution of rhodamine-B labeled CSP and the green channel (488 nm excitation with a 525/50 filter) shows the constitutive *gfp* reporter. (B) Magnified stereogram of the dotted region in (A).

**Supplementary Figure S5:**
Fluorescence of rhodamine B labeled CSP in planktonic cultures of *gfp*-expressing *S. mutans* and in growth medium without *S. mutans*.

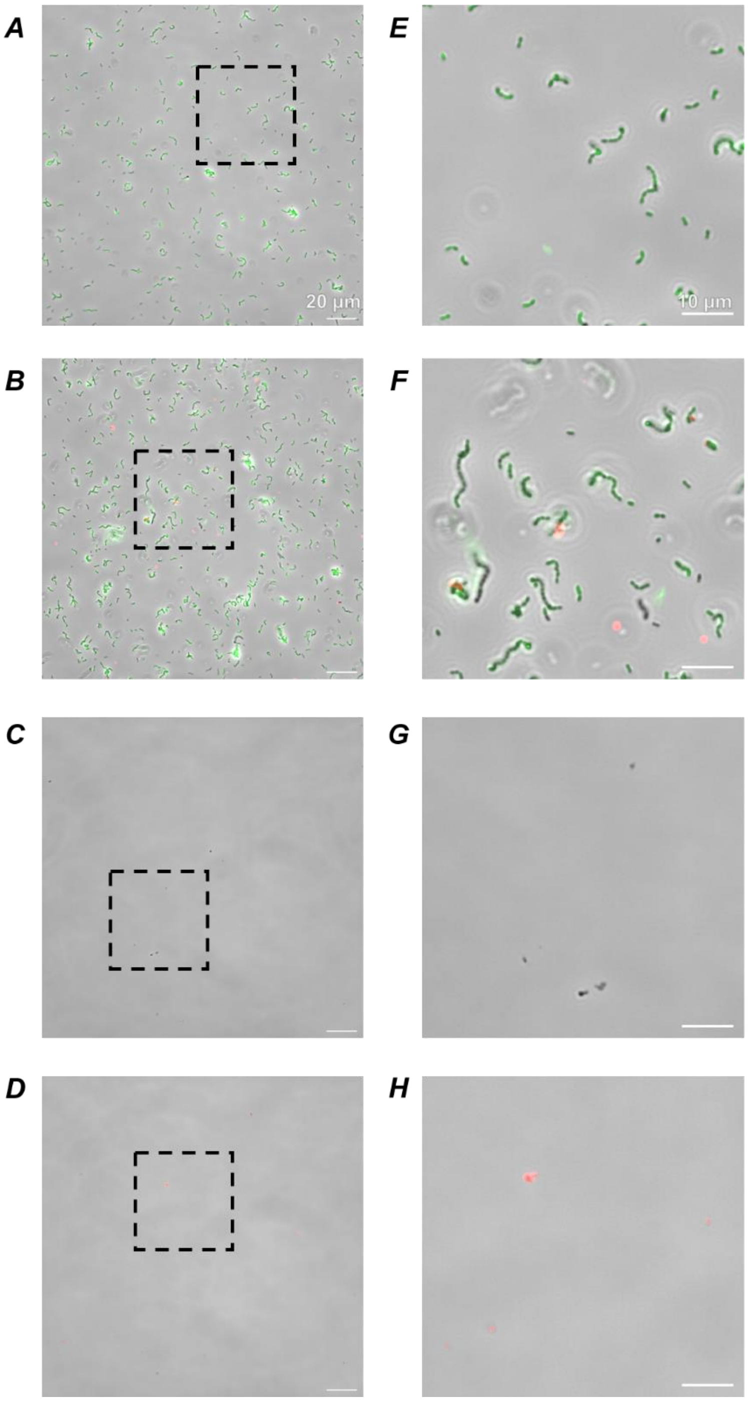

Phase contrast and fluorescence microscopy images of *gfp*-expressing *S. mutans* cells grown as a planktonic culture in BHI to an $OD_{600}$ of 0.2 before being washed and resuspended into fresh BHI with (A) 1 µM CSP or (B) 1 µM rhodamine B labeled CSP. Green channel shows the *gfp* reporter in the constitutive *Pldh-gfp* strain. Red channel shows the rhodamine B labeled CSP. (C) 5 µM CSP or (D) 5 µM rhodamine B labeled CSP in BHI with no *S. mutans* cells and 1 mg/ml spectinomycin. (E)-(H) Magnified views of the dotted regions in (A)-(D) respectively. Data shown are representative of 3 separate images collected for each condition.

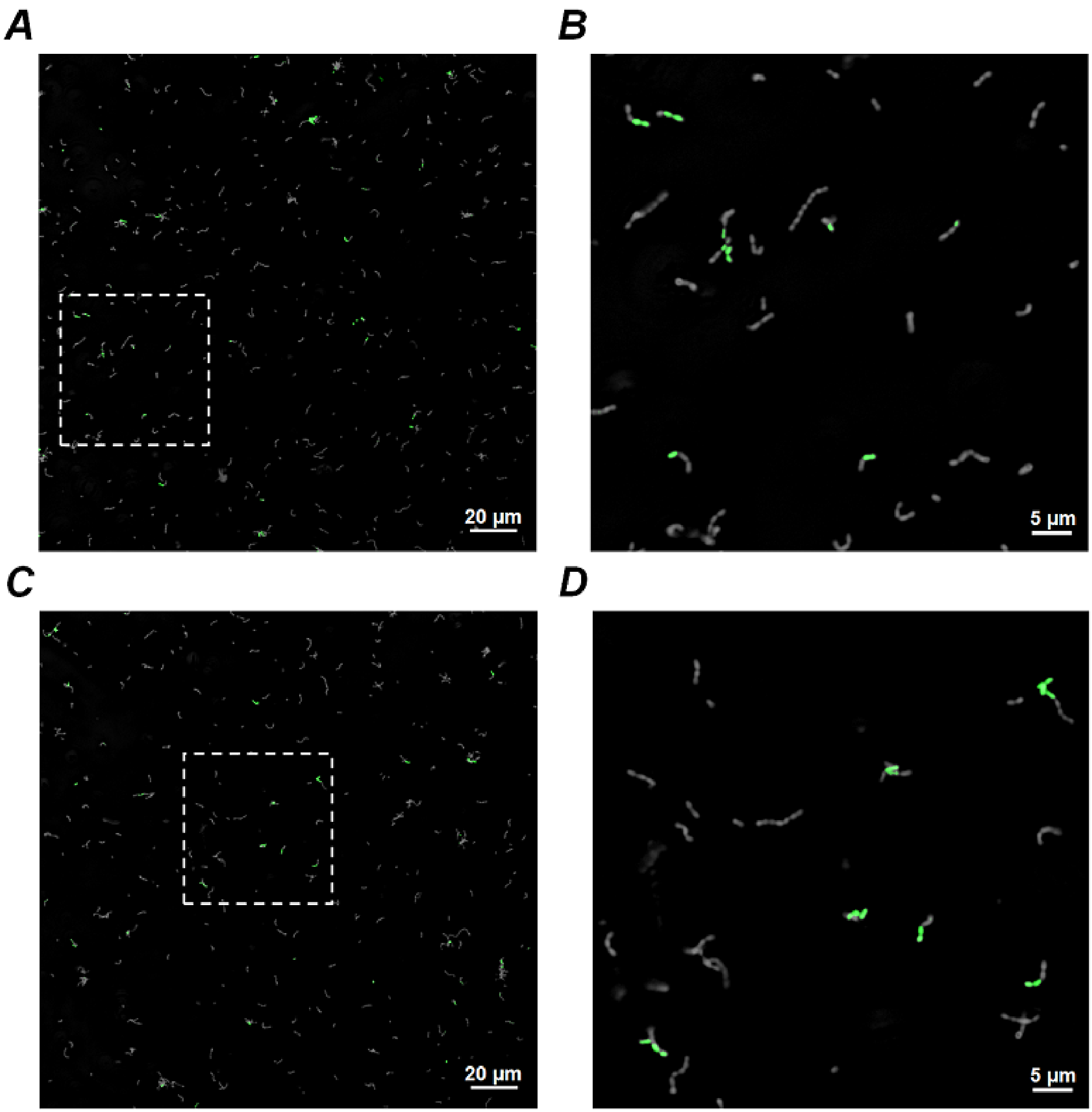

**Supplementary Figure S6**: Activation of the P*comX-gfp* reporter by 1 µM rhodamine B labeled CSP and by 1 µM unlabeled CSP

Microscopy images of P*comX-gfp* reporter strain grown planktonically and imaged in fluorescence and phase contrast. Culture was grown to $OD_{600}$ = 0.2 before being resuspended into fresh BHI medium with (A) 1 µM CSP or (C) 1 µM rhodamine B labeled CSP. Cells were incubated with CSP for approximately 2 h prior to imaging. (B)-(D) Magnified sections of the dotted regions in (A), (C) respectively. The phase contrast image is inverted (light on dark) and overlaid with the (green) fluorescence image to better reveal the activated cells. Data shown are representative of 3 separate images collected for each condition.

#### Correlation of P*comX* expression in sister cells of planktonic cultures

*Method*

For studies of planktonic *S. mutans*, overnight-grown cultures were washed twice and then diluted 25-fold into fresh BHI. After reaching $OD_{600nm}$ = 0.2, each culture was sonicated at 20% amplitude (Fisher FB120) for 20 seconds to break apart the cell chains into single cells. The culture was then resuspended into fresh BHI containing 1 µM of either CSP or (rhodamine B)-CSP. Subsequently, 4 µl of the culture was periodically extracted and dispersed on a glass coverslip and imaged in phase contrast and green fluorescence using a Nikon TE2000U inverted microscope equipped with a Photometrics Prime camera and a Nikon C-FL GFP HC HISN zero shift filter cube.

*Result*

Supplemental Figures S6 (A) and S6 (B) show cells 2 hours after addition of CSP. Cells chains have begun to form, and a subpopulation of roughly ~10% of cells respond to the CSP signal by expressing GFP. Notably, the fluorescence of adjacent cells on each growing chain is highly correlated: although the majority of cells show no fluorescence, most fluorescent cells are immediately adjacent to at least one fluorescent sister cell. These data demonstrate that sensitivity of P*comX* to exogenously provided CSP is heterogeneously distributed in the population overall, but is very highly correlated among sister cells. The data indicate that the "bowtie" doublet cells visible in Figure 1, which have a very similar appearance to the fluorescent chains in Supplemental Figure S4, are quite likely to be sister cells inheriting the CSP sensitivity of a common parent, rather than cells communicating by a diffusible signal. Supplemental Figure S6 (C) and (D) show a similar experiment in which cells are provided the rhodamine-B labeled CSP.

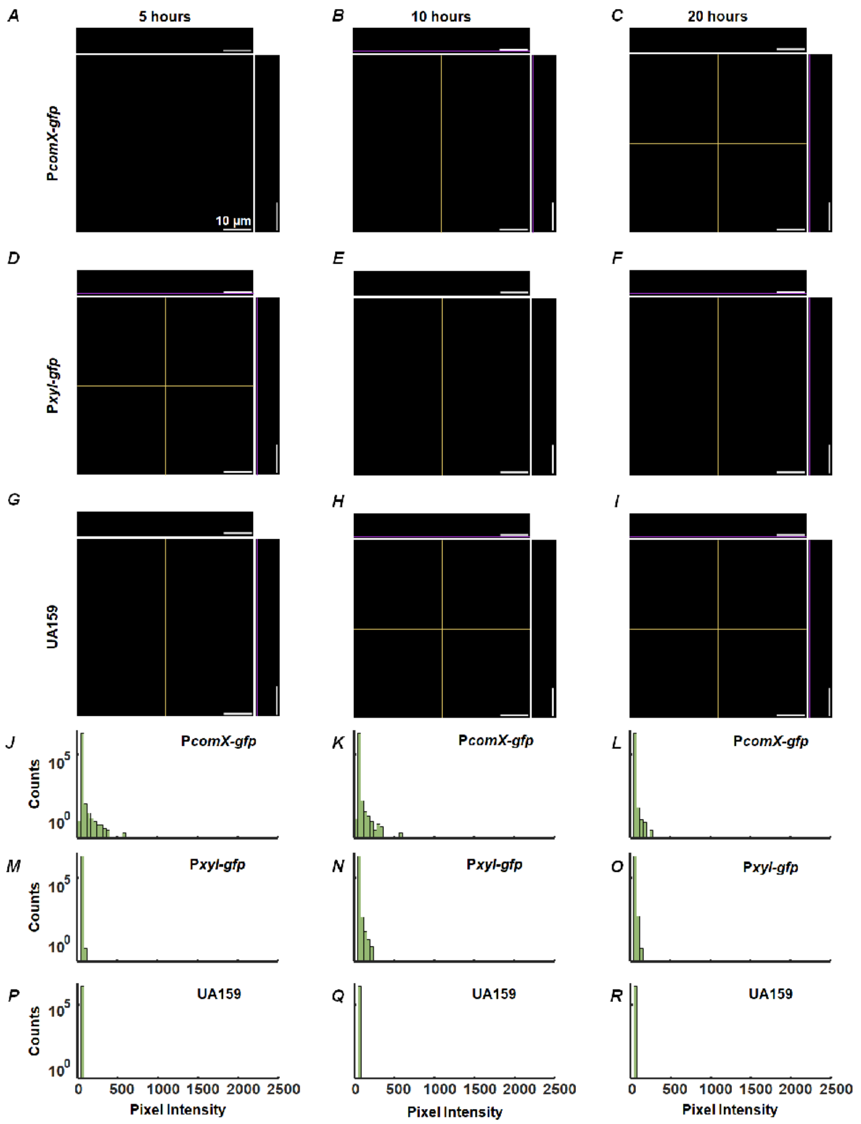

**Supplementary Figure S7**: Green fluorescence controls: biofilms of *PcomX-gfp* (without XIP), P*xyl-gfp* (without xylose), and UA159 background strains.

Multiphoton laser confocal (920 nm excitation) images of green fluorescence (525/25 nm detection) of biofilms grown to different time points. P*comX-gfp* biofilm was grown in defined media for (A) 5 h, (B) 10 h and (C) 20 h before incubation for an additional 3 hours in fresh FMC lacking XIP; P*xyl-gfp* strain was grown in defined media for (D) 5 h, (E) 10 h and (F) 20 h and incubated as above, without added xylose; (G)-(I) Background (non-reporting UA159) strain was grown by same procedure as (A)-(F). To match the brightness scales in Figure 6, the maximum pixel brightness is 2500 in (A)-(C) and (G)-(I), and 1500 in (D)-(F). The main panel within each image subpanel is a 2D *x*-*y* plane. The location of this plane within the depth *z* of the biofilm is indicated by the purple lines on the adjacent upper and right-side panels which are *x*-*z* and *y*-*z* sections respectively. The yellow lines on each central panel mark the *x* and *y* sections that are illustrated in the upper and right side panels. The total depth spans 30 µm; (J)–(R) Histograms of image brightness for above images. Histograms were generated by first binning each entire image stack, for (A)–(I), into volumes of 4x4x1=16 cubic pixels = 0.04 µm$^3$, as in Figure 6. Data shown are representative of 3 separate image stacks collected for each condition.

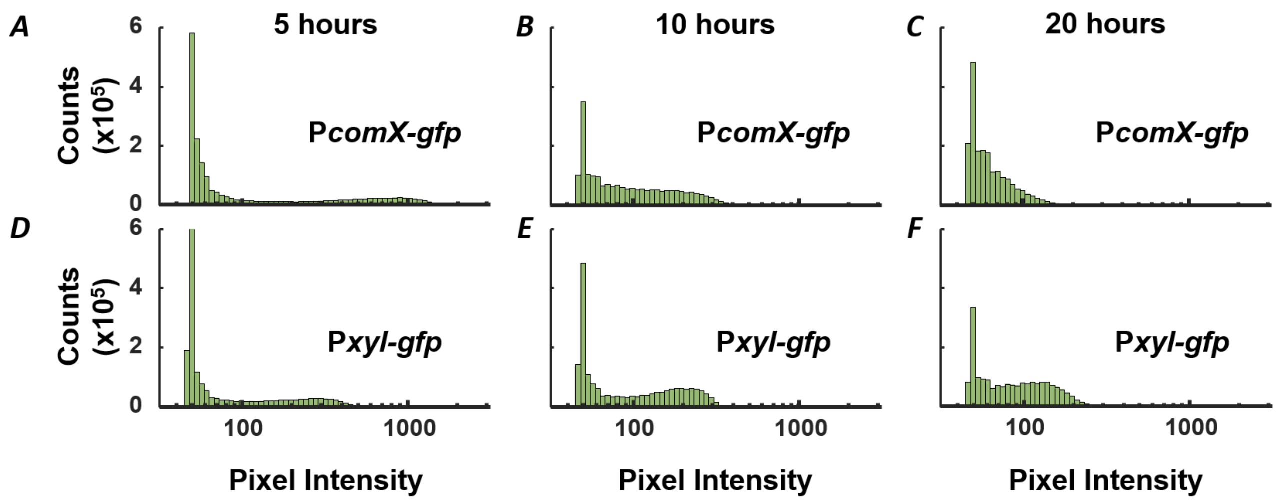

**Supplementary Figure S8**: Histograms of green fluorescence in biofilms of the *PcomX-gfp* (+XIP) and P*xyl-gfp* (+xylose) strains.

(A)-(F) Brightness intensity histograms for the image stacks of the P*comX-gfp* and P*xyl-gfp* biofilms in Figures 6A – F, respectively. Each entire image stack was first binned into volumes of 4x4x1=16 cubic pixels = 0.04 $\mu m^3$, as in Figure 6. The brightness histograms above were then constructed using a linear vertical scale with logarithmic bin spacing along the intensity axis. With linear binning (Figure 6) the fluorescence histograms are unimodal. However the logarithmic binning introduces a spurious bimodality (a second peak at intensity > 100) in all histograms except (C).
